# Mucin-degrading gut bacteria promote anti-parasitic immunity

**DOI:** 10.1101/2022.02.28.482289

**Authors:** Mathis Wolter, Marie Boudaud, Erica T. Grant, Amy Parrish, Alessandro De Sciscio, Seona Thompson, Jean-Jacques Gerardy, Michel Mittelbronn, David J. Thornton, Andrew J. Macpherson, Richard K. Grencis, Mahesh S. Desai

## Abstract

**BACKGROUND & AIMS:** Host-secreted gastrointestinal mucus plays a key role in the expulsion of intestinal nematode parasites. A balance between mucin secretion by the host and the gut microbial mucin foraging is essential to maintain the intestinal homeostasis, yet little is known about how changes in the mucin–microbiome interactions affect worm infections. Here, we aimed to examine how mucin foraging activity by the microbiome changes the course of parasitic worm infections by modulating the host immune responses.

**METHODS:** We utilized a gnotobiotic mouse model containing a synthetic human gut microbiota that allows for: 1) a complete removal of the mucin-degrading bacteria from the community; and 2) diet-driven manipulation of the microbiota toward mucin foraging. We infected mice with a murine nematode, *Trichuris muris*, which resembles human infection with *Trichuris trichiura*. We examined the temporal dynamics of worm infection including worm burden and the host immune responses, and coupled these readouts to the microbial changes and mucin foraging activity.

**RESULTS:** The absence of mucin-degrading bacteria in the microbiota enhances susceptibility to parasitic infection—evidenced by higher worm number—by promoting stronger Th1 immune responses. Dietary fiber deprivation increases the microbial mucin-foraging activity, which coincides with a shift in host immune responses from susceptible (chronic, Th1 type) to resistant (acute, Th2 type), thereby promoting worm clearance.

**CONCLUSIONS:** Our results provide mechanistic insights into how the colonic mucin-degrading bacteria promote anti-parasitic immunity through modulation of the host immune responses. Our study documents a clinically-relevant, novel link in the microbiome–parasite–host immune axis that is useful prerequisite knowledge in managing parasitic infections.

## Introduction

Intestinal parasitic worms (helminths) make an invaluable model to study the host immunomodulatory mechanisms due to their ability to subvert the host immune responses^1^. In this context, the murine parasite *Trichuris muris—*a relevant biological model for the closely related human parasite *Trichuris trichiura—*has been extensively investigated^2^. *T. muris* is an especially attractive experimental model as either a susceptible or resistant immune response can be investigated^2,3^. Susceptibility in mice, which is more reflective of the naturally-occurring chronic infections^4^, is generally characterized by a Th1 response resulting in long-term worm infections and chronic colitis. On the other hand, resistance is characterized by an acute Th2 response leading to worm clearance and tissue repair^2^.

Since they occupy the same host environment, it is no surprise that enteric parasites such as helminths strongly interact with the gut microbiome^5,6^. Although these interactions may lead to the suppression of either the gut microbiome or the helminth^5–7^, in case of *T. muris*, bacterial–helminth crosstalk is essential for worm hatching, as the parasite eggs fail to hatch and establish in germ-free environments^8–10^. Once the worm has settled in the intestinal epithelium, the bacterial–helminth dynamics appear mainly unidirectional, with helminths altering the microbiome composition through various mechanisms, including through helminth excretory-secretory products or host anti-microbial peptides, released in response to the parasite^11–15^. Furthermore, the helminths alter the nutrients available to the microbiome, indirectly influencing bacteria selection^16^. The effect of the infection on host immunology has an additional impact on the microbiome, as the worms or their metabolic byproducts can alter toll-like receptor (TLR) expression profiles^17,18^ or responsiveness of host immune cells to TLR ligands^19^ and interfere with local IgA-production^17^. While the gastrointestinal (GI) mucins are essential for worm clearance^20–22^ and the impacts of the helminth infection on the microbiome during worm clearance are documented at the broader level of microbial phylogeny^23–28^, questions remain regarding how the mucus–microbiome interactions are affected by and affect worm infection and the underlying host immune responses.

Worm expulsion during an acute response is largely driven by goblet-cell secreted gastrointestinal mucin that is tightly linked to the Th2-type immune response^22^. Along these lines, mice lacking the *Muc2* gene, which encodes the major glycoprotein of the GI mucus, show a delayed clearance of *T. muris*^29^ and *Muc5ac* deficient mice completely fail to clear the helminth^22^.

We have previously shown that a fiber-deprived microbiota leads to a disrupted balance towards excessive gut microbial mucin foraging activity^30–32^. As the balance between mucin secretion and its consumption by the gut microbes is critical for intestinal homeostasis^33,34^, here, we hypothesized that excessive mucin foraging by the microbiota alters the course of the parasite infection by modulating the immune response. To test this hypothesis, we employed a tractable 14-member synthetic human microbiota (14SM) characterized for the capability of individual members to degrade various glycans including mucin oligosaccharides^31,35^. In a longitudinal experimental setup, using ex-germ-free mice with a mucin-foraging or a non-mucin-foraging synthetic microbiota, we studied how mucin–gut microbiome interactions shape anti-parasitic immunity. Our findings suggest a mechanism of how mucin-degrading bacteria promote anti-parasitic immunity, which sheds a new light on the microbiome– parasite–host immune interactions.

## Methods

### Ethical statement

All animal experiments were performed according to the “Règlement Grand-Ducal du 11 janvier 2013 relatif à la protection des animaux utilisés à des fins scientifiques” based on Directive 2010/63/EU on the protection of animals used for scientific purposes, and approved by the Animal Experimentation Ethics Committee of the University of Luxembourg and by the Luxembourgish Ministry of Agriculture, Viticulture, and Rural Development (national authorization no. LUPA2020/02). The mice were housed in accordance with the recommendations stated by the Federation of European Laboratory Animal Science Associations (FELASA).

### Experimental design and dietary treatment

8–23 weeks old C57BL/6 mice were housed in ISOcages with up to five animals per cage. Sterile food and water were provided *ad libitum* and light cycles consisted of 12 hours. Before the administration of 14SM, the germ-free status of the mice was confirmed using aerobic and anaerobic culturing of stools samples. Mice were intragastrically gavaged with 0.2 ml of the 14SM or 10SM gavage mix as described previously^35^. Before and for 6–9 days following the gavage, mice were maintained on a standard mouse chow, which we refer to as Fiber-rich (FR) diet. Afterwards, half of the mice were assigned to a Fiber-free (FF) diet and the rest remained on the FR diet. Twenty days after the diet switch, the mice were infected by oral gavage with a high dose of approximately 450 eggs of *T. muris* strain E and observed for up to 30 days post-infection (dpi). Mice were euthanized by cervical dislocation to determine worm counts and other final readouts. A piece (∼0.5 cm) of the cecum was fixed in methacarn in order to determine goblet cell numbers. The remaining part of the cecum and 2 cm of the attached colon were frozen at –20 °C to determine *T. muris* worm burden. A piece (∼0.5 cm) of the colon was emptied from content, longitudinally cut-opened and weighted before an incubation of 24 h at 37 °C, 5 % CO_2_, in culture medium (RPMI with L-glutamine, 5% fetal bovine serum, 100U penicillin-streptomycin) for analysis of cytokine secretion. The remaining colonic tissue and mesenteric lymph nodes were stored in RNAprotect Tissue Reagent (Qiagen, Hilden, Germany; catalog no. 76106, according to the manufacturer’s protocol) or processed for immune cell isolation and FACS analysis. Serum of the euthanized mice was collected and stored at –20 °C.

### Parasitological techniques

*T. muris* maintenance and infection was performed as previously described^36^. Mice were orally infected with ∼450 eggs and worm burdens were assessed by counting the number of worms present in the cecum and 2 cm of the attached colon on days 20, 25 and 30 post infection, as described previously^36^. The researcher, S.T., who counted the worms, was blinded for both the individual time points and dietary groups.

### ELISA

Lipocalin-2 ELISA was performed on stool samples using the lipocalin-2/NGAL DuoSet ELISA R&D system (Bio-Techne, Minneapolis, MN, USA; catalog no. DY1857) according to the manufacturer’s instructions and samples were prepared as previously described^31^. MCPT1 concentration of serum samples was determined using the MCPT-1 (mMCP-1) Mouse Uncoated ELISA Kit from Invitrogen (Waltham, MA, United States, catalog. no. 88-7503-88) according to manufacturer’s instructions. Serum ELISA for total IgG1 and IgE were performed by coating plates using 0.5 µg/µl rat α mouse IgE purified UNLB (Imtec Diagnostics, 1130-01) or rat α-mouse IgG1 purified UNLB (Imtec Diagnostics, Ardmore, OK, United States; catalog no. 1144-01) capture antibody diluted in 0.05 M carbonate/bicarbonate buffer (pH 9.6). Washing steps were performed using 1% Tween, 154 mM NaCl, 10 mM Trizma Base.

Blocking of the plates was performed using 1% w/v of BSA in a TBS buffer (15 mM Trizma-acetate, 136 mM NaCl & 2 mM KCl). Mouse IgG1 Isotype Control UNLB (Imtec Diagnostics, Ardmore, OK, United States; catalog no 0102-01) or Mouse IgE Isotype Control (UNLB), Southern Biotech (Imtec Diagnostics, Ardmore, OK, United States; catalog no 0114-01) served as controls. Serum samples were two-fold diluted using the aforementioned TBS buffer with 0.1% w/v Tween-20 and 1% BSA. The secondary antibody was Goat anti-mouse IgG1-AP, Southern Biotech (Imtec diagnostics, Ardmore, OK, United States; catalog no 1071-04) or Goat anti-mouse IgE-AP, Southern Biotech (Imtec diagnostics, Ardmore, OK, United States; catalog no 1110-04), respectively. The substrate solution consisting of 0.5 mg/ml phosphate substrate (Sigma-Aldrich, St. Louis, MO, United States; catalog no. S0642-200 TAB) in 1 mM AMP and 0.1 mM MgCl_2_•6H_2_O and plates were read at 405 nm using a SpectraMax ABS PLUS spectrophotometer (Molecular Devices, San Jose, CA, United States). A detailed protocol can be found in the supplementary methods section. Parasite specific IgG1 and IgG2c ELISA was performed as previously described^37^.

### LEGENDplex bead-based immunoassay

The LEGENDplex assay (BioLegend, San Diego, CA, USA) was performed on serum samples or colon tissue supernatant according to manufacturers’ instruction.

### RNA extraction and RT-qPCR from mesenteric lymph nodes, colonic tissue

A detailed protocol can be found in the supplemental methods section.

### RNA-Seq

RNA extracted from samples at 30 dpi were utilized for performing RNA-Seq. Illumina Stranded Total RNA Prep with Ribo-Zero Plus was used to prepare the RNA sequencing library according to the reference guide’s instructions. Sequencing was performed using NovaSeq 6000 SP Reagent Kit v1.5 (Illumina, San Diego, CA, USA) on an Illumina NovaSeq 6000 system.

### Flow cytometry analysis of lamina propria immune cells

Colonic immune cells were isolated using a Lamina Propria Dissociation Kit (Miltenyi Biotec, Bergisch Gladbach, Germany) on the GentleMACS Dissociator (Miltenyi Biotec), following manufacturer’s instructions. Cells were washed, counted, and stained as previously described^38^. Samples were acquired on a NovoCyte Quanteon flow cytometer (ACEA Biosciences Inc., San Diego, CA, USA). Raw fcs files were analyzed in FlowJo version 10.8.1 as previously described^38^.

### Statistical analyses

Statistical analysis was performed using Prism 9.2.0. (GraphPad Software, Inc., San Diego, CA, USA). Error bars in all figures represent SEM. Unless specified in the figure legend, a two-way analysis of variance (ANOVA) was performed in Prism and *p* values were adjusted with the original FDR method of Benjamini and Hochberg. P values lower than 0.1 are reported in the figures.

## Results

### Dietary fibers tune the worm infection dynamics

We colonized 22-weeks-old germ-free, C57BL/6 mice with a 14-member synthetic human microbiota (14SM), fed them a fiber-rich (FR) or a fiber-free (FF) diet, and then infected them with a high dose of *T. muris* consisting of approximately 450 eggs from the same egg suspension **(Fig. 1A)**. Throughout the experiment, none of the mice exhibited any obvious physiological abnormalities. While FF-fed mice were overall heavier compared to the FR-fed mice, the weights of both groups remained largely stable over time **(Fig. S1A)**. In line with our published work^31,39^, the FF diet allowed proliferation of two dominant mucin-degrading bacteria *Akkermansia muciniphila* and *Bacteroides caccae* and reduction of two prominent fiber-degrading bacteria *Eubacterium rectale* and *Bacteroides ovatus* **(Fig. 1B)**. Overall, colonization with the synthetic human microbiota permitted establishment of a low-level *T. muris* infection. Interestingly, although the worm burdens were statistically similar between FR and FF groups on 20 and 25 dpi, subsequent infection dynamics significantly differed between both groups, with FF-fed mice clearing the pathogen by 30 dpi and FR-fed mice remaining chronically infected **(Fig. 1C)**.

**Figure 1.**
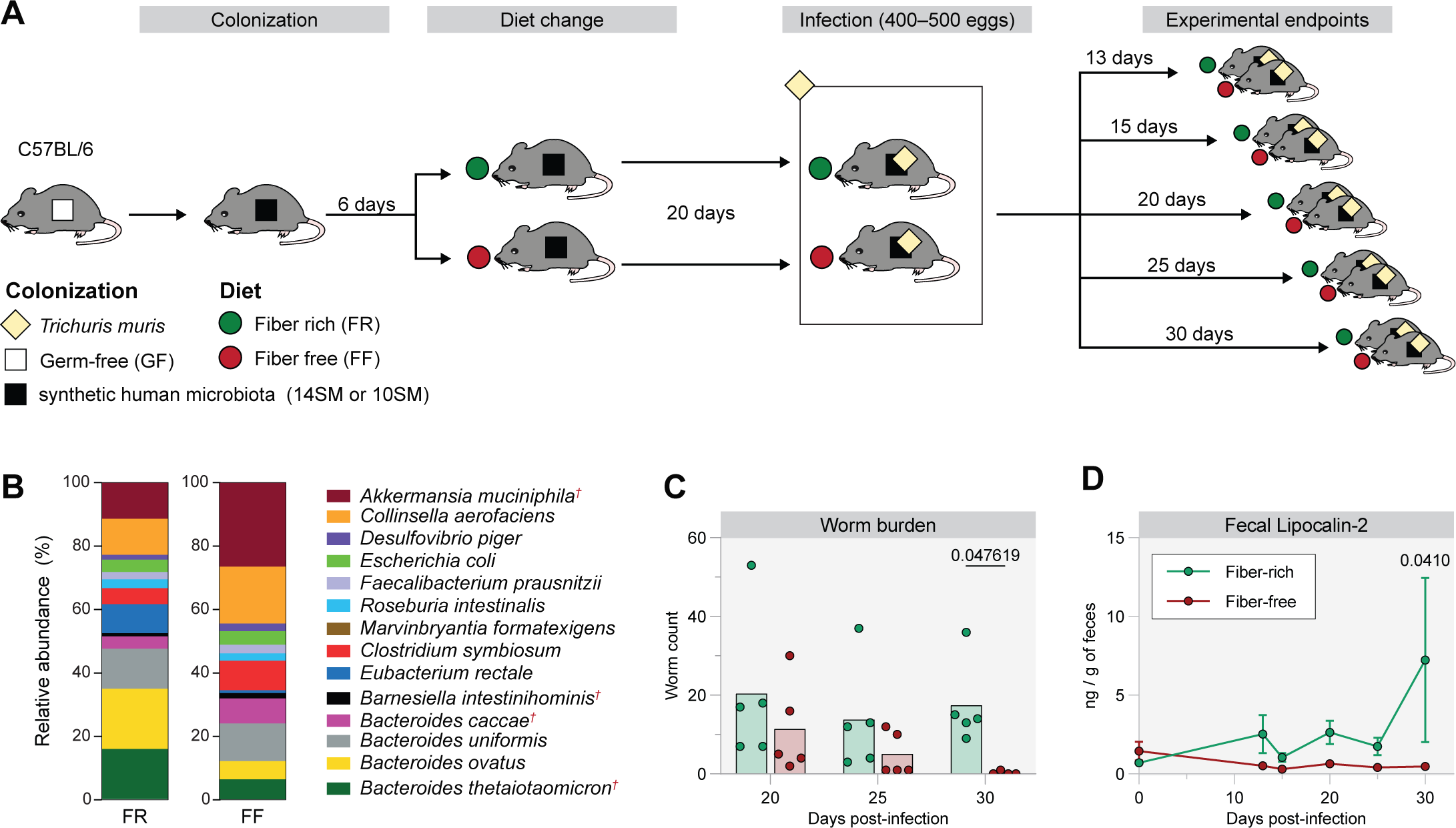
Diet drives changes in mucin-degrading bacteria expansion and alters worm infection. (A) Experimental timeline. (B) Relative bacterial abundance at the time of infection (day 0), determined by qPCR on fecal DNA. Red latin crosses denote known mucin-degrading bacteria. (C) Worm burdens assessed at 20, 25 and 30 dpi. Each dot represents a mouse (n=4–5). Multiple Mann-Whitney test, *p* values adjusted with the FDR method of Benjamini and Hochberg. (D) Fecal Lipocalin-2 (LCN-2) levels assessed by ELISA before (day 0) and after infection with *T. muris* (13, 15, 20, 23, 30 dpi) (n=4–5). *p* values indicate comparisons between FR and FF groups.

In order to investigate whether the two distinct diets and/or the skewed gut microbiome composition toward mucin polymer degradation caused an altered baseline inflammation prior to infection with *T*. *muris*, we determined fecal lipocalin-2 (LCN-2) levels, which serves as a neutrophilic marker for low-grade inflammation^40^. There were no significant differences between FR and FF groups at day 0, however, after infection, the levels of LCN-2 were significantly higher in the FR group **(Fig. 1D)**.

### Diet-driven *T. muris* chronic infection or clearance are supported by Th1 or Th2 immune responses

Chronic *T. muris* infection has been associated with Th1 inflammation while clearance of the worm is usually attributed to a strong Th2 response^2^. To better understand the change in host immunological responses that drive differential *T*. *muris* infection dynamics, we analyzed the host colonic gene expression at 30 dpi **(Fig. 2A, B, Fig. S1B, C and Table S1)**. We found 147 and 184 genes upregulated in the FR- and FF-fed mice, respectively **(Fig. 2A and Tables S2, S3)**. Ingenuity Pathway Analysis (IPA) revealed that both the interferon signaling and Th1 pathways were upregulated in the FR-fed mice **(Fig. 2B)**, which was consistent with the failure to successfully clear the parasite among FR-fed mice **(Fig. 1C)**. In accordance with the upregulation of the interferon signaling and Th1 pathways, related disease pathways, such as osteoarthritis, systemic lupus erythematosus and even neuroinflammation, were also upregulated as their gene expression profiles overlap with those of a Th1 response **(Fig. 2B)**. Similarly, the Th1 response is potentially linked to the observed increase in antimicrobial responses that results in the activation of cytotoxic T cells, lymphopoiesis, increased apoptosis of macrophages and a reduced replication of viruses **(Fig. S1B)**. In contrast to this, the increased MSP-RON signaling in macrophage pathway hints at a Th2 response under FF conditions **(Fig. 2B),** as an activation of the MSP-RON pathway causes M2 macrophage polarization^41^, which is associated with a Th2-type immune response against *T. muris*^42,43^. Furthermore, the upregulation of the white adipose tissue browning pathway is also in line with this observation, as M2 macrophage differentiation has been linked to adaptive thermogenesis^44^.

**Figure 2.**
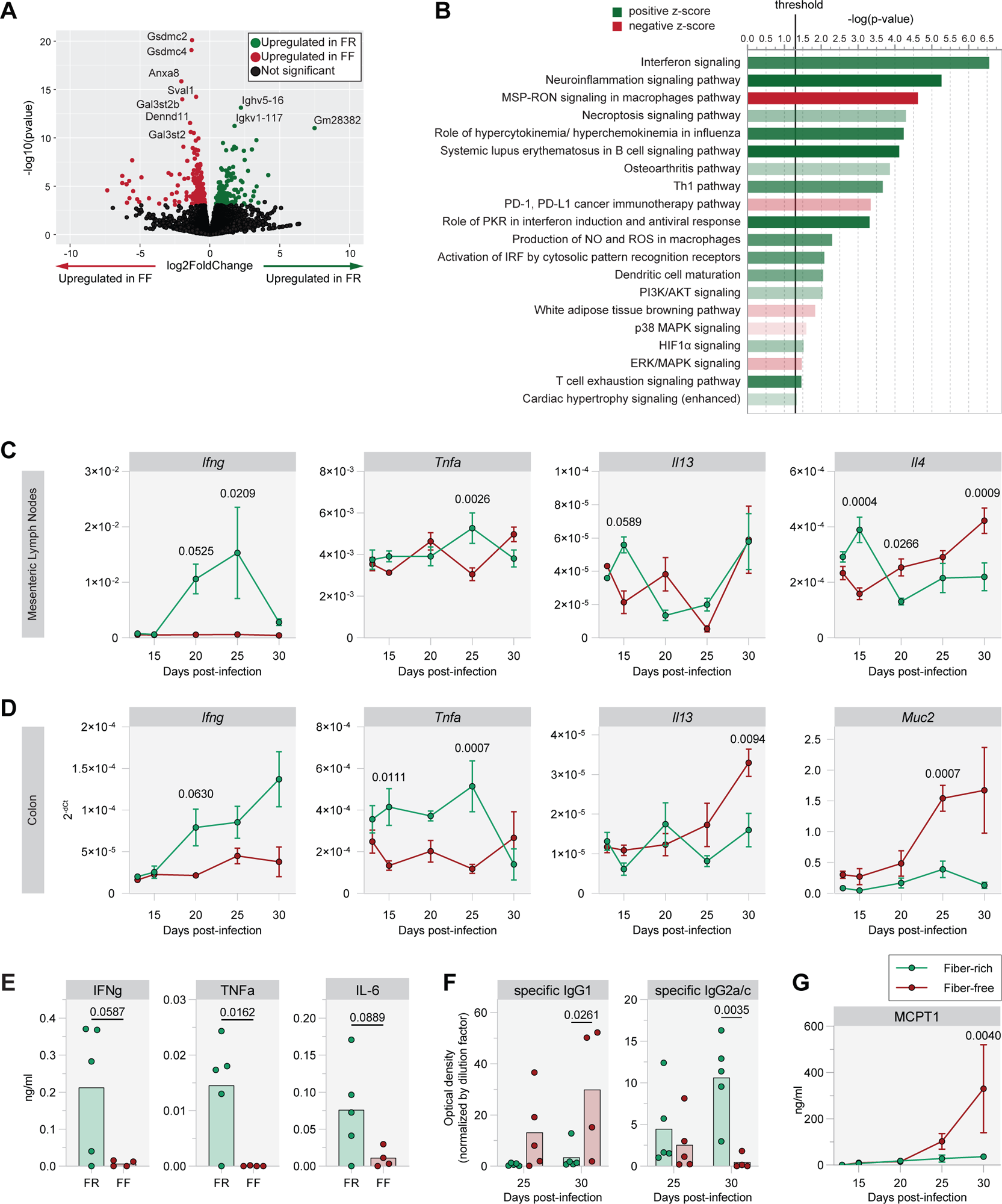
Dietary fibers skew the anti-parasitic immune response towards Th1-type responses. (A, B) Analysis of data generated by RNA-seq of RNA extracted from colonic tissue at 30 dpi (n=4). (A) Volcano plot of the distribution of all differentially expressed genes. (B) Barplot showing significantly up- or downregulated host pathways ((Fiber Rich (FR)/Fiber Free (FF)) determined by Ingenuity Pathway Analysis (IPA). The higher the opacity of the bar, the more up- or downregulated the pathway. (C, D) Relative expression of indicated transcripts in (C) mesenteric lymph nodes and (D) colonic tissue over time after *T. muris* infection (n=3–5); *p* values indicate comparisons between FR and FF groups. (E) Serum cytokine concentrations at 30 dpi (n=4–5). Unpaired, two-tailed t-test. (F) Parasite-specific IgG1 and IgG2a/c concentrations in mouse serum (n=4–5). (G) Serum mast cell protease 1 (MCPT1) concentrations (n=2–5); *p* values indicate comparisons between FR and FF groups.

In order to further support conclusions from our RNA-Seq data, we used RT-qPCR to investigate transcript levels of cytokines in mesenteric lymph nodes (MLNs) and colon **(Fig. 2C, D, Table S4)**. In accordance with the global transcriptome analysis **(Fig. 2A, B)**, the FR-fed mice exhibited a Th1-type response in the MLN, characterized by a peak of *Ifng*, *Tnfa* and *Il1b* transcripts at 25 dpi and a continuously increasing expression of the Th1 transcription factor *Tbet* **(Fig. 2C and Fig. S1D)**. This Th1-type response was also observed in the colon with an increasing expression of *Ifng* over time and a peak of *Tnfa* expression at 25 dpi **(Fig 2D)**, as well as higher titers of IFNɤ, TNFα and IL-6 in the serum of FR-fed mice at 30 dpi **(Fig. 2E)**. Despite an initial induction of the anti-parasitic Th2-type response in the MLN of FR-fed mice at 15 dpi, *Il13* and *Il4* transcript levels decreased by 20 dpi, consistent with the simultaneous increased expression of the counteracting Th1-type cytokines **(Fig. 2C)**. By contrast, transcript levels of the Th2-type cytokines IL-4, IL-5 and IL-13 and the Th2 transcription factor GATA-3 were upregulated over time in either the MLN or colons of FF-fed mice **(Fig. 2D and S1D)**. Further supporting the efficient anti-parasitic response in FF-fed mice, the transcript levels of *Muc2*, the major component of the intestinal mucus, was strongly upregulated in FF-fed mice by 25 and 30 dpi, while remaining low in FR-fed mice **(Fig. 2D)**.

Assessment of serum concentrations of total IgE and IgG1, both of which are reflective of the protective Th2-type responses against helminths, resulted in no significant changes, albeit an increased trend was observed for both antibodies at 30 dpi **(Fig. S1E)**. We then assessed parasite-specific IgG1 and IgG2, characteristic of Th2 and Th1 responses, respectively^45^. Chronic *T. muris* infections are usually characterized by significantly elevated serum levels of parasite-specific IgG2a/c and a low increase in parasite-specific IgG1^45^. In contrast, protection against *T. muris* is characterized by high parasite-specific IgG1 levels, but a lack of IgG2a/c^45^. In accordance with the observed worm burdens **(Fig. 1C)**, at 30 dpi, we saw an increase in specific IgG1 in the serum of mice fed the FF diet, while we saw an increase in specific IgG2a/c in mice fed the FR diet **(Fig. 2F)**. Furthermore, serum MCPT1—a proxy for IgE-induced mast cell degranulation^46^—increased at 30 dpi in the FF-fed group **(Fig. 2G)**. These data are in line with an increased tendency of total IgE in the FF-fed group at 30 dpi, and further support the elevated anti-helminth type 2 immunity under fiber deprivation.

Overall, our data support that worm infection in mice hosting the 14SM and being fed fibers, induce a Th1-type immune response that is driven by an overall increase in IFNγ. On the other hand, in absence of dietary fibers, a similar infectious dose of *T.muris*, induces a Th2-type immune response, resulting in the successful clearance of the worm. We have previously shown that fiber deprivation leads to excessive gut microbial mucin foraging activity^30–32^. Since mucus secretion is essential to the anti-parasitic immunity, and as the gut microbiome is intimately connected to changes in the diet and the host immune responses, we hypothesized that mucin-degrading bacteria promote anti-parasitic immunity.

### Mucin-degrading bacteria and absence of fiber promote anti-parasitic immunity

To study the causal mechanism of mucin-degrading bacteria in modulating immunity to helminth infection, we next infected 8-to-12-weeks-old mice, both male and female, colonized with a 10-member synthetic microbiota (10SM) consisting of the 14SM minus the 4 previously characterized mucin-degrading bacteria^31^. In this experiment, 14SM-colonized mice had lower worm burdens compared to the previous experiment (maximum 19 worms versus 53, respectively) (**Fig. 1C and 3A**), and no difference was detected between males and females (data not shown). However, in line with the previous experiment (**Fig. 1C**), the mice fed the FR diet had a higher worm burden than mice on the FF diet which completely cleared the worms by 30 dpi (**Fig. 3A**). The 10SM FR-fed mice harbored worms at all three time points, whereas FF-fed mice presented with few or no worms at any time point (**Fig. 3A**). Nevertheless, all mice developed parasite-specific IgG, confirming that they were properly infected (**Fig. 3B**). Interestingly, mice fed the FR diet and devoid of mucin-degrading bacteria (10SM FR) had a higher worm burden by 30 dpi compared to 14SM-colonized FR mice (**Fig. 3A**), while they were similar between both groups on 20 and 25 dpi (**Fig. S2A)**. Additionally, among both FF groups at 30 dpi, 10SM-colonized mice had a higher titer of serum specific IgG1 (**Fig. 3B).** On the other hand, among both FR groups at 30 dpi, 10SM-colonized mice had higher amounts of fecal lipocalin-2 **(Fig. S2B**). These data suggest that mucin-degrading bacteria limit worm infection by altering anti-parasitic immunity.

**Figure 3.**
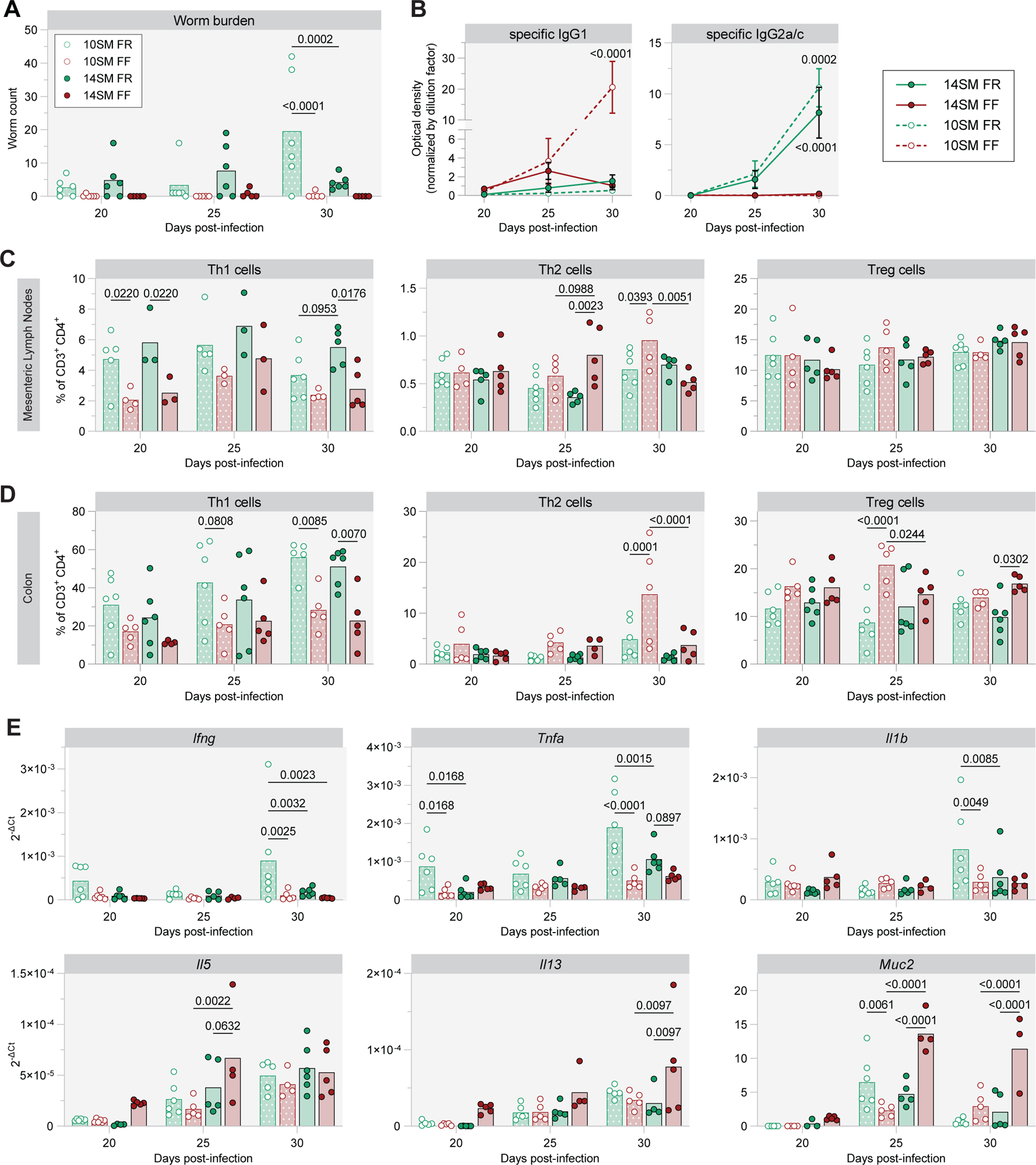
Mucin-degrading bacteria promote the anti-parasitic immunity. (A) Worm burdens assessed at 20, 25 and 30 dpi in 14SM- and 10SM-colonized mice fed a fiber-rich (FR) or a fiber-free (FF) diet (n=5–6). (B) Parasite-specific IgG1 and IgG2a/c concentrations in mouse serum (n=5–6). *P* values indicate comparisons between FR and FF groups among 10SM- or 14SM-colonized mice. (C, D) Proportion of indicated immune cell populations in (C) the mesenteric lymph nodes and (D) colonic tissue (n=3–6). (E) Relative expression of indicated transcripts in colonic tissue (n=3–6).

To better assess the anti-parasitic immune response, we analyzed the immune profiles of mesenteric lymph nodes (MLN) and colons at 20, 25 and 30 dpi. Regardless of bacterial colonization, flow cytometry analysis of immune cell populations in the MLN and colons revealed a higher proportion of Th1 cells in FR-fed mice, in contrast to more Th2 (both MLN and colon) and Treg (colon only) cells in the FF-fed mice (**Fig. 3C, D and S2C**). This diet effect was further supported by transcript and protein expression profiles of Th1-related cytokines (IFNγ, TNFα and IL-1β) in the colons of FR-fed mice and more Th2-related cytokines (IL4, IL-5, IL-9 and IL-13) and MCPT1 in the colons of FF-fed mice (**Fig. 3E and S3**).

Among FR-fed mice, 14SM-colonized mice had lower expression levels of Th1-related cytokine transcripts *Ifng*, *Tnfa* and *Il1b* than 10SM-colonized mice at 30 dpi (**Fig. 3E**). Although these trends were not always consistent at the protein level (**Fig. S3B**), they are perfectly in line with the higher worm burden in 10SM FR mice (**Fig. 3A**), corroborating that the presence of mucin-degrading bacteria further modulates intensity of the anti-parasitic immune response. Among FF-fed mice, 14SM colonization tended to increase the proportion of Th2 cells in the MLN at 25 dpi but then reduced it at 30 dpi in both the MLN and colon (**Fig. 3C, D**). Supporting an earlier T cell response in 14SM-colonized mice, the colonic transcript levels of the growth factor IL-2 and the Type 2 promoting cytokines IL-25 and IL-33 were already higher at 20 dpi in FF-fed, 14SM-colonized mice compared to FF-fed, 10SM-colonized mice (**Fig. S3A**).

Intriguingly, FF-fed, 10SM-colonized mice had increased proportions of Tregs, ILC1 and ILC3 in the colon at 25 dpi (**Fig. 3D and S2C**), which may impair a timely Th2-driven anti-parasitic response^47^. Further confirming an impaired anti-parasitic response in FF-fed, 10SM-colonized mice, the increased proportion of colonic Th2 and ILC2 at 30 dpi was poorly reflected by IL-5 and IL-13 cytokine production (**Fig. 3D, S2C and S3B**). Consistently, *Muc2* transcript was strongly increased in FF-fed, 14SM-colonized mice but not in 10SM-colonized mice (**Fig. 3E**). Additionally, at 25 dpi, 14SM-colonized FF-fed colons also produced more IL-10 and Th17-related cytokines (IL17-A, IL-17F and IL-22) compared to all other groups (**Fig. S3B**), suggesting a more efficient barrier healing and protective process in the presence of mucin-degrading bacteria and in the absence of dietary fibers. These results strongly support that mucin-degrading bacteria and absence of fiber act together to promote an anti-parasitic immune response.

### Mucin-degrading bacteria promote mucus production under fiber deprivation

As a result of a strong type-2 immune response, mucus production is a key mechanism of host defense during parasite infection and can be regulated at the transcriptional, translational and post-translational levels. From 15 to 30 dpi, both FR- and FF-fed 14SM-colonized mice had moderately increased numbers of goblet cells per crypt, but no difference was seen between dietary groups (**Fig. 4A**). Transcriptomic analysis of colon tissues at 30 dpi revealed that the expression of numerous genes involved in mucus production and glycosylation pathways were upregulated in FF-fed mice compared to FR-fed mice (**Fig. 4B**). Moreover, targeted qPCR revealed that, in addition to *Muc2* (**Fig. 2C and 3E**), *Muc5ac* expression was higher in 14SM-colonized FF-fed mice at 30 dpi, compared to FR-fed mice or FF-fed, 10SM-colonized mice (**Fig. 4C**). Similarly, the combination of both mucin-degrading bacteria and fiber deprivation induced the expression of *C1galt1*, *B3gnt6*, *Fut2*, *Galnt3* and *Gal3st2*, which encode key enzymes involved in mucin glycosylation (**Fig. 4C, Table S4**). These results suggest that mucin-degrading bacteria support production of mucus under fiber deprivation, and possibly ensure its proper glycosylation so that this new mucus can be secreted in order to expulse the worms.

**Figure 4.**
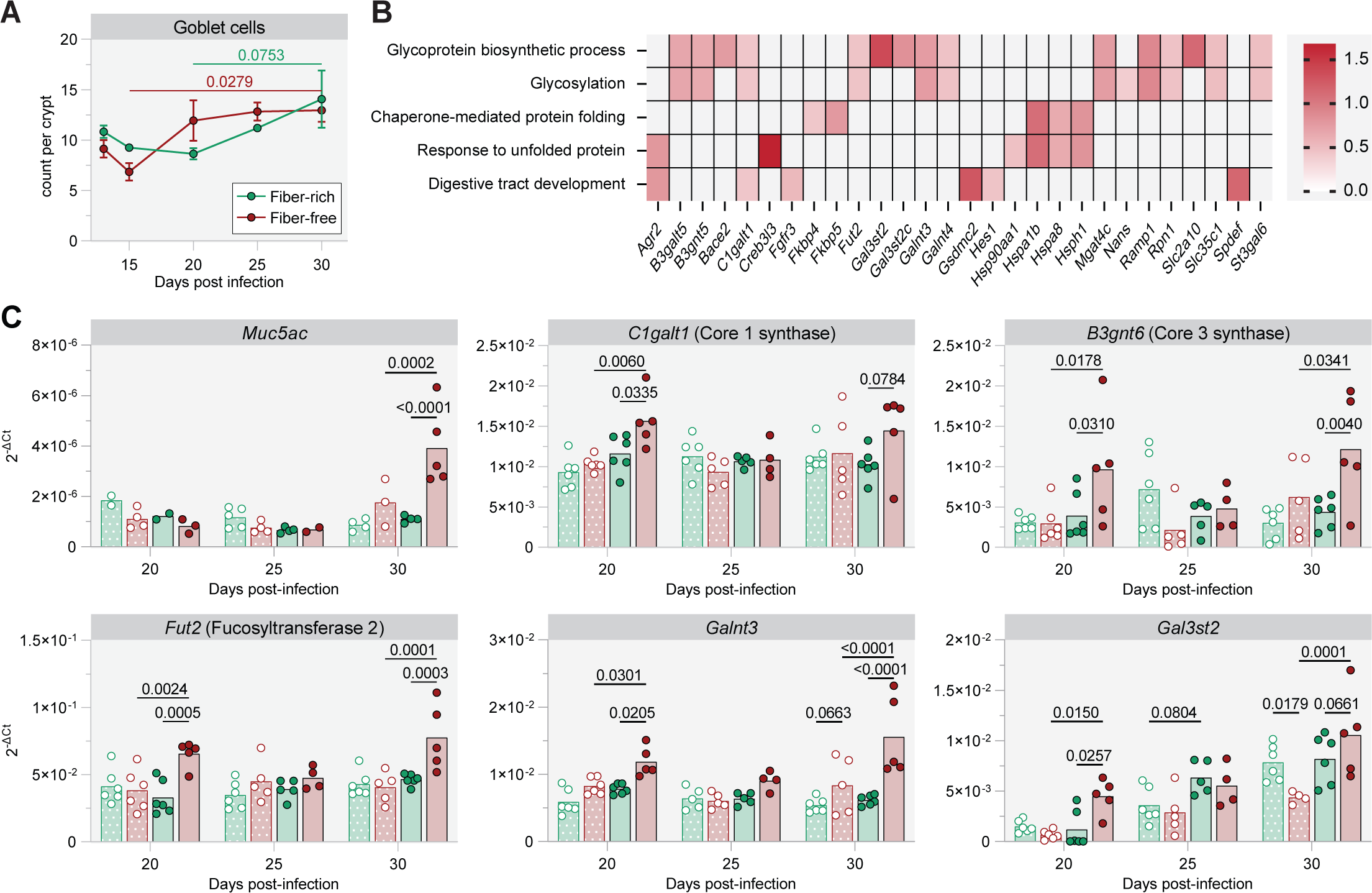
Mucin-degrading bacteria promote mucus production under fiber deprivation. (A) Goblet cell counts per crypt of *T. muris* infected mice determined by histological assessment of methacarn-fixed cecum sections (n=2–5). (B) Relative gene expression (FF/FR) at 30 dpi of the 5 most significant activated pathways in the colonic tissue under FF conditions (n=4 per group). Pathways are based on GO terms for biological processes and REVIGO was used to reduce redundant terms. (C) Relative expression of indicated transcripts in colonic tissue (n=2–6).

### Increased mucin–microbiome interactions promote worm clearance

To better understand the role of specific microbiota members in the course of the parasite infection, we first analyzed the composition of the fecal microbiota (**Fig. S4**). Surprisingly and despite the reproducible phenotypes showed above, the 14SM microbiota composition at the time of infection was different between the first experiment (**Figs. 1, 2**) and the second experiment (**Fig. 3**) (**Fig. S4, 5**, **Supplementary results**). The analysis of the relative abundance over time for individual bacteria revealed different profiles between groups with little reproducibility between independent experiments, except for *A. muciniphila* and *B. caccae* (**Fig. 5A, B**, **S4B and S6, 7**). Despite different initial relative abundances between the two experiments, these 2 mucin degraders expanded significantly and reproducibly from 14 to 20 dpi (**Fig. 5A, B**), a timepoint when mice usually start to expel their worms.

**Figure 5.**
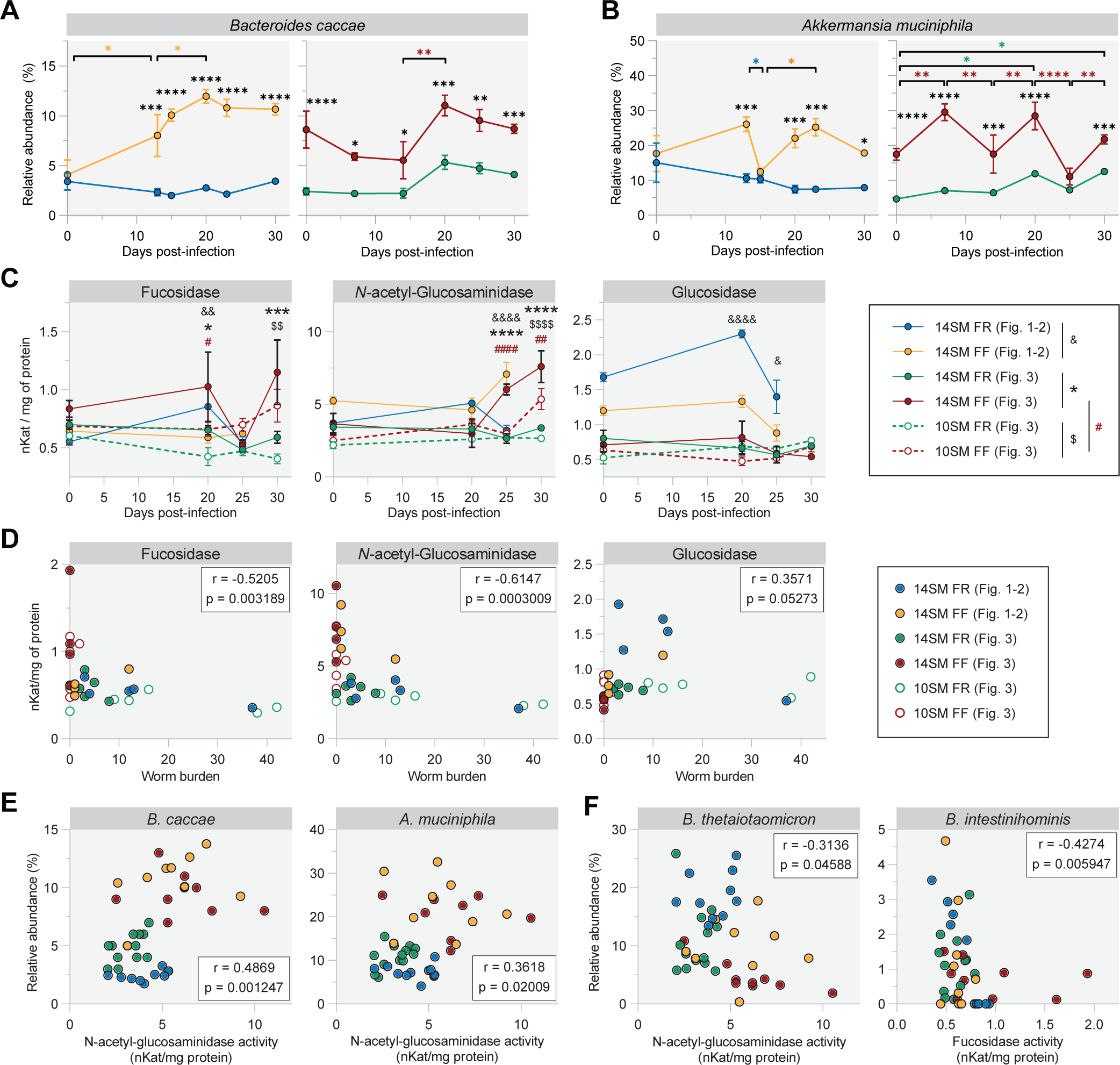
Increased mucin–microbiome interactions promote worm clearance. (A, B) Relative abundance of *B. caccae* (A) and *A. muciniphila* (B) over the course of *T. muris* infection in 14SM-colonized mice (n=2–6). Significance labels indicate comparisons between FR versus FF groups (black), or between indicated timepoints among FF-fed mice of Fig. 1–2 (yellow), FR-fed mice of Fig. 3 (green) and FF-fed mice of Fig. 3 (red). (C) Glycan-degrading enzyme activities of the gut microbiome determined by stool-based p-nitrophenyl glycoside-based enzyme assays. α-fucosidase and β-N-acetyl-glucosaminidase are key mucin-degrading enzymes, while β-glucosidase serves as a control for general glycan-degrading activity (n=3–6). Significance labels indicate comparisons between ^&^FR versus FF groups among 14SM-colonized mice from Figs. 1, 2, *FR versus FF groups among 14SM-colonized mice from Fig. 3, ^$^FR versus FF groups 10SM-colonized mice from Fig.3, and ^#^10SM versus 14SM groups among FF-fed mice. ^&^/*/^#^p<0.05; ^&&^/^$$^/^##^p<0.01; ***p<0.001; ^&&&&^/****/^$$$$^/^####^p<0.0001. (D, E) Spearman correlations between indicated bacteria relative abundance and (D) N-acetyl-glucosaminidase activity (n = 41–70) or (E) fucosidase activity (n = 40). (F) Spearman correlations between worm burdens and enzymatic activities (n=31).

Consistent with the evolving mucus production and microbiota composition, the enzymatic activities targeting either mucin glycans only (fucosidase and *N*-acetyl-glucosaminidase) or both mucin and dietary glycans (glucosidase) evolved over time during *T. muris* infection (**Fig. 5C, Tables S5–7**). The activities of mucin-targeting enzymes were higher in 14SM-colonized and FF-fed mice (**Fig. 5C**), and correlated negatively with the worm burden (**Fig. 5D**), further supporting a role for mucin-degrading bacteria in facilitating worm expulsion. Of note, fucosidase activity in 14SM-colonized FF-fed mice (**Fig. 5C**) followed the expression profile of the fucosyltransferase transcript *Fut2* (**Fig. 4C**), decreasing at 25 dpi to increase again by 30 dpi, supporting a microbial enzyme activity that may be driven by the glycosylation profile of the mucins. Finally, among the four mucin-degrading strains, the relative abundance of *B. caccae* and *A. muciniphila* correlated positively with *N*-acetyl-glucosaminidase activity (**Fig. 5E and Table S6**), while the relative abundance of *B. thetaiotaomicron* and *B. intestinihominis* correlated negatively with *N*-acetyl-glucosaminidase and fucosidase activities, respectively (**Fig. 5F**, **Tables S5, 6**).

## Discussion

Helminth infection models have been profoundly utilized to elucidate the underlying host responses for the past several decades, yet open questions remain about the mechanisms of how the intricate interactions between the gut microbiome and GI mucus affect worm infection dynamics including the host immune responses. Here, we shed insight into the role of bacterial mucin foraging in altering the host immune responses during the infection with *T. muris*. Using a functionally characterized synthetic human microbiota that allows either an ablation of microbial mucin foraging or increase in microbial mucin foraging through reduced fiber consumption, our study supports a model in which the mucin-degrading bacteria promote anti-parasitic immunity, that can be further enhanced by reduced dietary fiber **(Fig. 6)**. Our results identify mucin-degrading bacteria as an innovative link in the gut microbiome–parasite axis, suggesting a causal mechanism through which the anti-parasitic immunity is a modulated.

**Figure 6.**
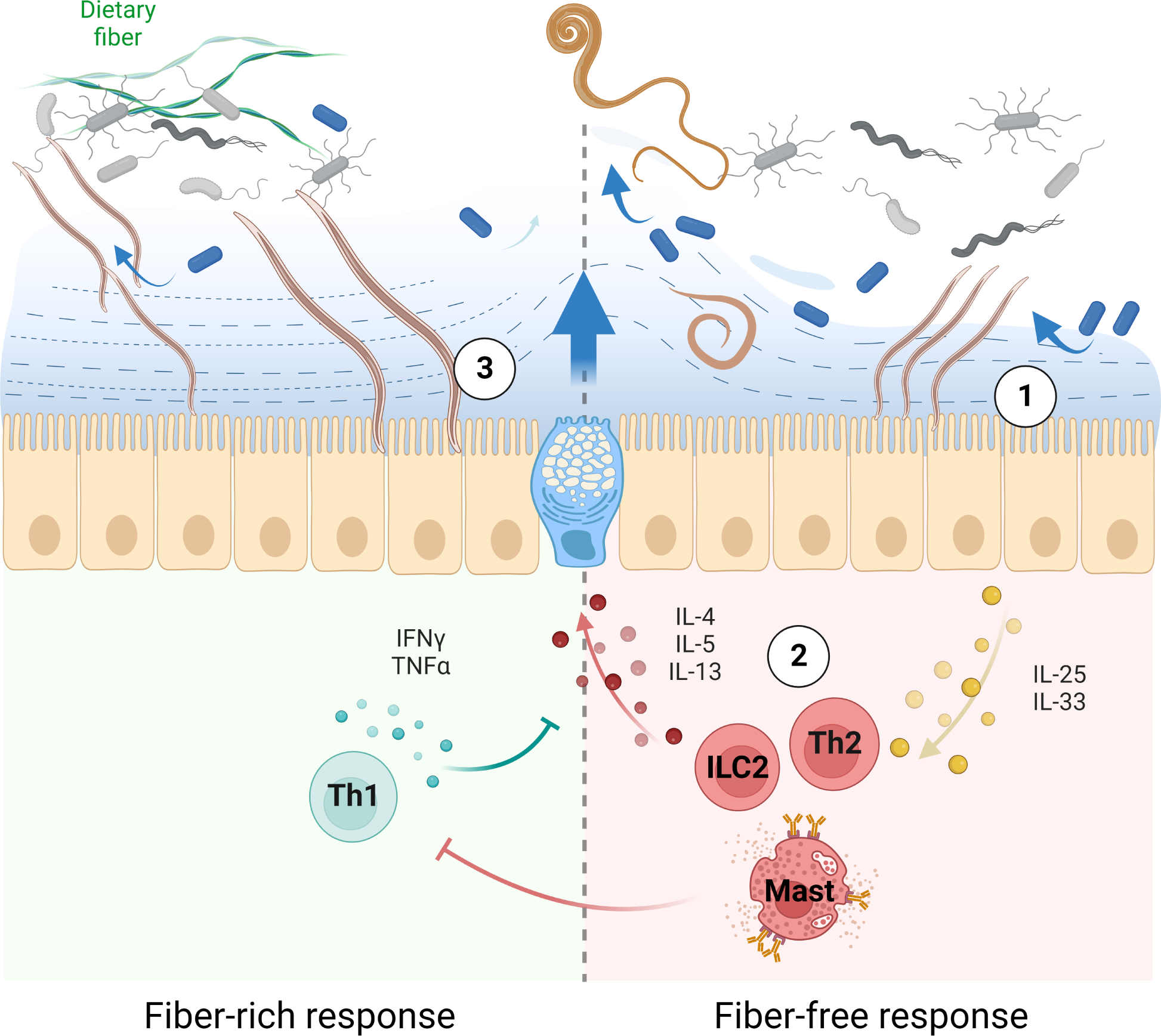
Model for the role of mucin-degrading bacteria in facilitating anti-parasitic immunity and *T. muris* clearance. (1) Fiber deprivation promotes microbial mucus foraging and thinning, which may facilitate hatching of the worm to the epithelium and its sensing by the host. (2) This sensing induces a strong type-2 response, leading to mucus production and preventing the worm to stay and develop. (3) Under fiber deprivation the worms reaching the epithelium promote an IFNg response, which inhibits the anti-parasitic response and allows the worm to settle and grow.

It is well known that bacteria are important for *T. muris* worm hatching^8,9^, although the specific bacterial members involved are not well defined and it is possible that the selected bacteria in our synthetic human microbiota might not be ideal for inducing egg hatching for mouse *Trichuris*, resulting in a lower level of infection of *T. muris* that provides a chronic low-level infection characteristic of infections under natural conditions. Thus, our tractable synthetic microbiota, that facilitates functional interpretations, is a good model to understand the mechanisms linking the worm infection and mucin–microbiome interactions, which would be difficult to decipher in such great detail using a complex, native microbiota. Although the fiber-deficient mice, lacking the mucin-degrading bacteria, contained none to very few worms, these mice still mounted a robust anti-parasitic immune response (**Fig. 3B**). Differences in the parasitic load observed between the two experiments (**Figs. 1, 2 vs. Fig. 3**) could be due to several factors such as the age of the mice used for these experiments (22 weeks vs. 8–12 weeks) or in some other unknown factors, yet both experiments provided similar key outcomes. We noticed inconsistency between some readouts e.g., immune cell profiles (MLNs and colons) and serum cytokines at protein level, which is not unexpected for such detailed, complex longitudinal data sets. Nevertheless, overall, the data from our gnotobiotic mouse models allow us to adequately support our main conclusions.

Our study shows an IFNγ-dominated Th1 response, along with a chronic *T. muris* infection, when mice were fed a normal chow (which we called a fiber-rich (FR) diet), in contrast to a Th2 response and worm clearance under fiber deprivation **(Figs. 1–3 and 6)**. Two previous studies^27,48^ observed a similar Th1 response when specific pathogen free mice—containing their native, complex microbiotas—were fed diets containing purified 10% inulin or dried chicory leaves. Nevertheless, these studies did not provide an explanation for the underlying mechanisms. Using our 14SM model, we offer an explanation that the mucin–microbiome interactions driven by dietary changes alter the host immune responses and the worm infection dynamics **(Fig. 6)**. Our study lays groundwork for investigating future molecular mechanisms of how mucolytic bacteria drive specific alterations in mucus glycosylation and how such alterations impact host immune response to worm infection.

*T. muris* has been shown to broadly alter the host’s gut microbiome composition^23–28^. Here, repeating the experiment in several batches of animals revealed different, most likely age-dependent, profiles of microbiota composition over the course of the infection (**Figs. S4–7**). Nevertheless, the infectious phenotype was reproducible as well as a decline of *A. muciniphila* at 15 dpi followed by an expansion of both *A. muciniphila* and *B. caccae* at 20 dpi in fiber-deprived mice (**Fig. 5A, B**). The role of mucin-degrading bacteria such as *A. muciniphila* and *B. caccae* in our model is evident because colonization with a synthetic microbiota without mucin-degrading bacteria prevented the fiber deprivation-induced Th2 immunity (**Fig. 3**).

In contrast to the observations in *Muc2^-/-^* mice^29^, in our model, increased microbial mucin foraging under fiber-deprived conditions did not delay the parasite clearance, but instead promoted the clearance. These results are not unexpected as genetic ablation of Muc2 largely destabilizes the mucus structure, whereas the increased microbial mucin foraging seems to either resist transition to a Th1 response and aids in a Th2 response. We could not detect an increase in goblet cell numbers/mucin production associated with this (**Fig. 4A**). Nevertheless, we observed increased transcription of genes involved in mucin glycosylation (**Fig. 4B, C**), which has been shown previously to alter susceptibility to *T. muris*^49^. Considering that mucin glycosylations are built-up by the host glycosyltransferases and targeted by the microbial enzymes^50^, our results suggest that fiber deprivation promotes changes in mucin glycosylation that facilitates mucus degradation by the fiber-deprived microbiota and aids in clearance of *T. muris*.

Given the high morbidity associated with the parasitic infections in humans^51^, our study has generated important prerequisite knowledge for clinical settings: 1) for treating parasitic infections – in the absence of co-infections, periodic fiber deprivation or fasting to boost microbial mucin foraging could be potentially employed in the clinic to manage worm infections; and 2) for harnessing the immunomodulatory properties of helminths for their potential therapeutic use in inflammatory disorders^52^ or allograft rejection^53^. Further research is required about how the model proposed in our study **(Fig. 6)** could be translated into the clinic.

## Correspondence

Mahesh S. Desai, PhD Luxembourg Institute of Health Department of Infection and Immunity 29, rue Henri Koch, L-4354 Esch-sur-Alzette Luxembourg Tel: +352 26970-389 Fax: +352 26970-390

## Author Contributions

M.S.D. supervised the study and obtained research funding; M.W., M. B., D.J.T., R.K.G. and M.S.D. conceived and designed the study; M.W., M. B., E.T.G., A.P., A.D.S., and S.T. performed the experiments; M.W., E.T.G., R.K.G., A.P. and M.B. analyzed the data; J.-J. G. and M.M. prepared slides for histology; A.J.M. supplied germ-free mice; M.W., M. B., and M.S.D. primarily wrote the manuscript. All authors reviewed and approved the manuscript.

## Data Transparency Statement

All data, study materials and analytical methods will be made available to other researchers upon request.

## Supporting information

Supplementary methods, supplementary results and supplementary figure/table legends

Supplementary Tables combined

---

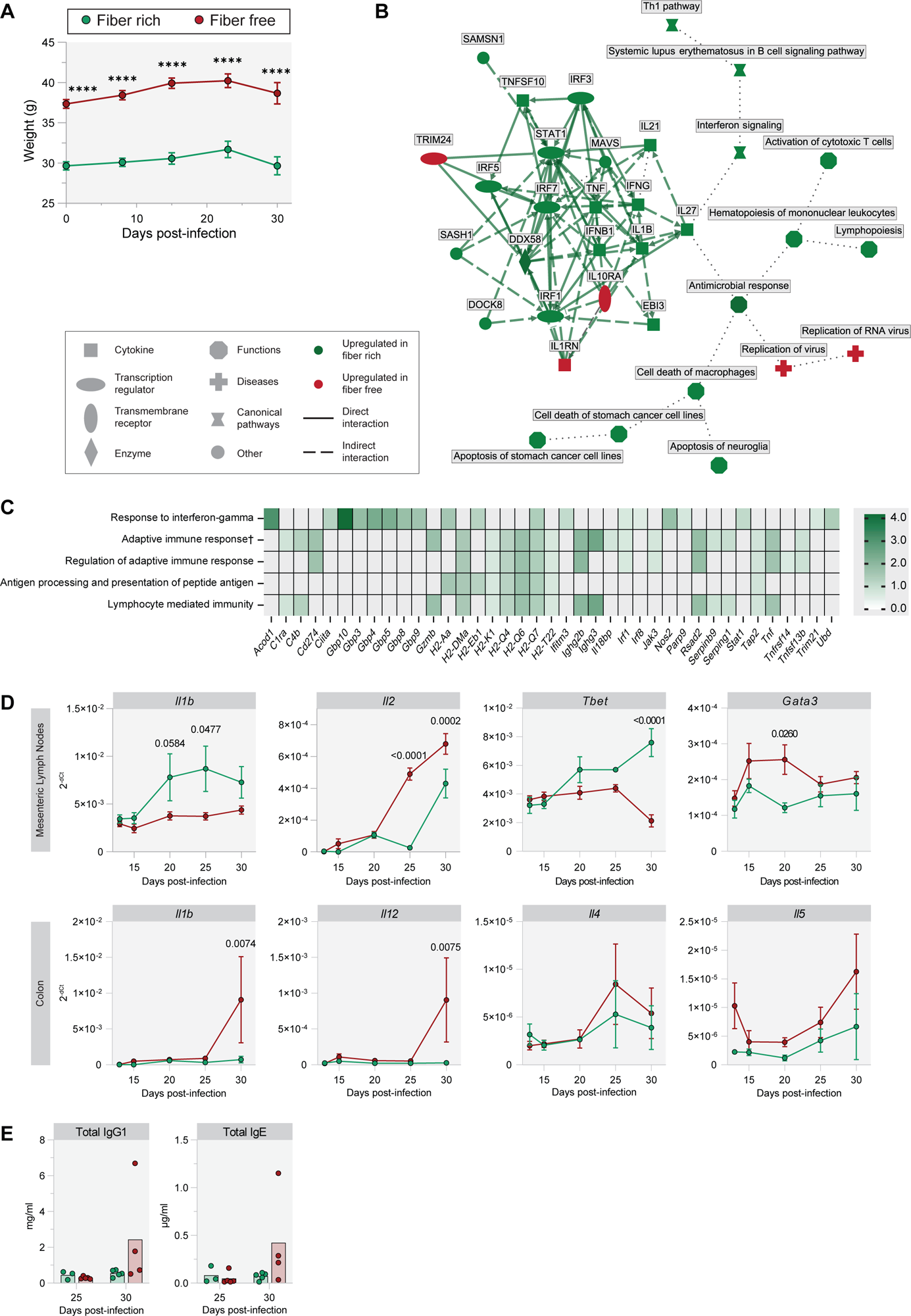

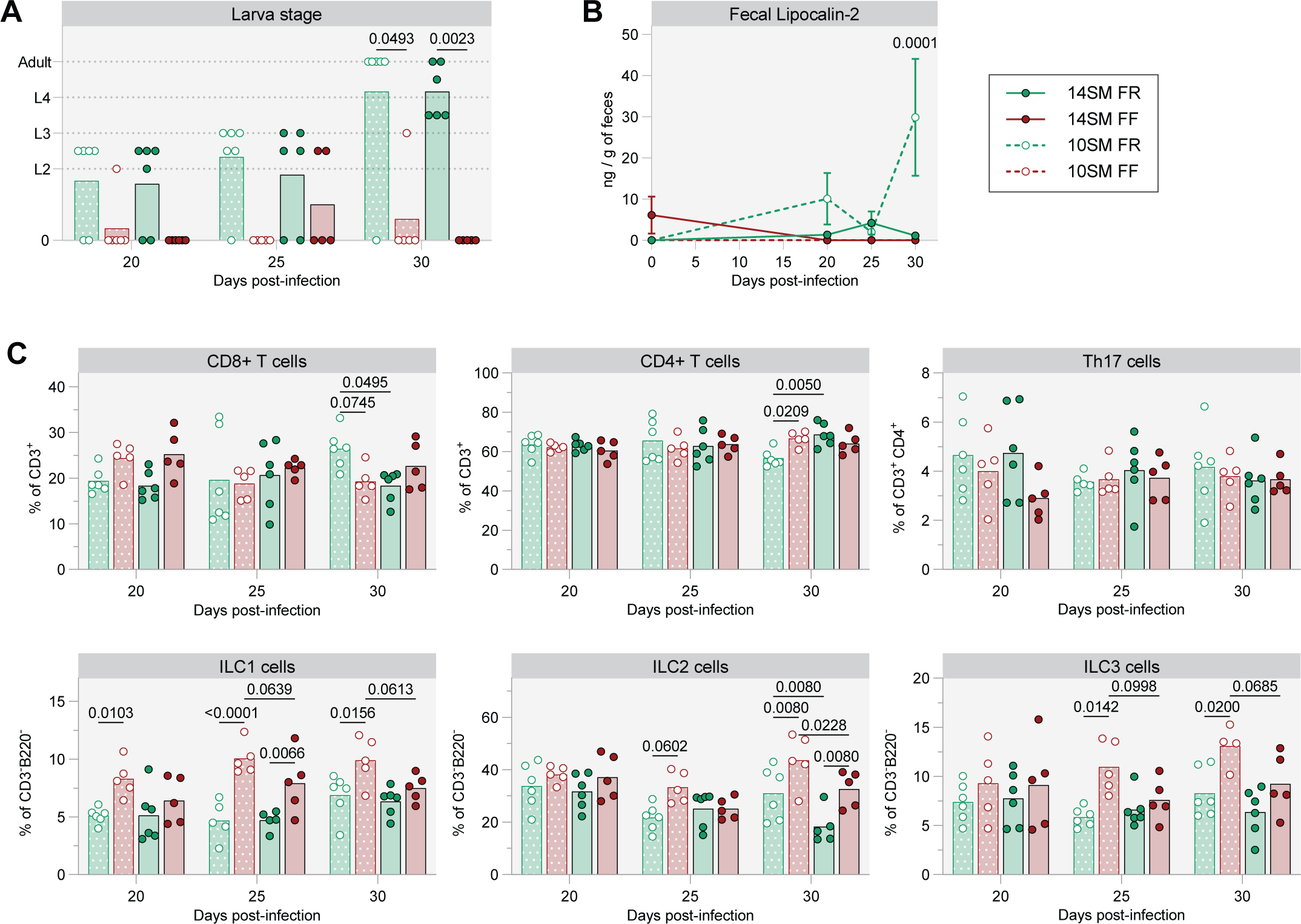

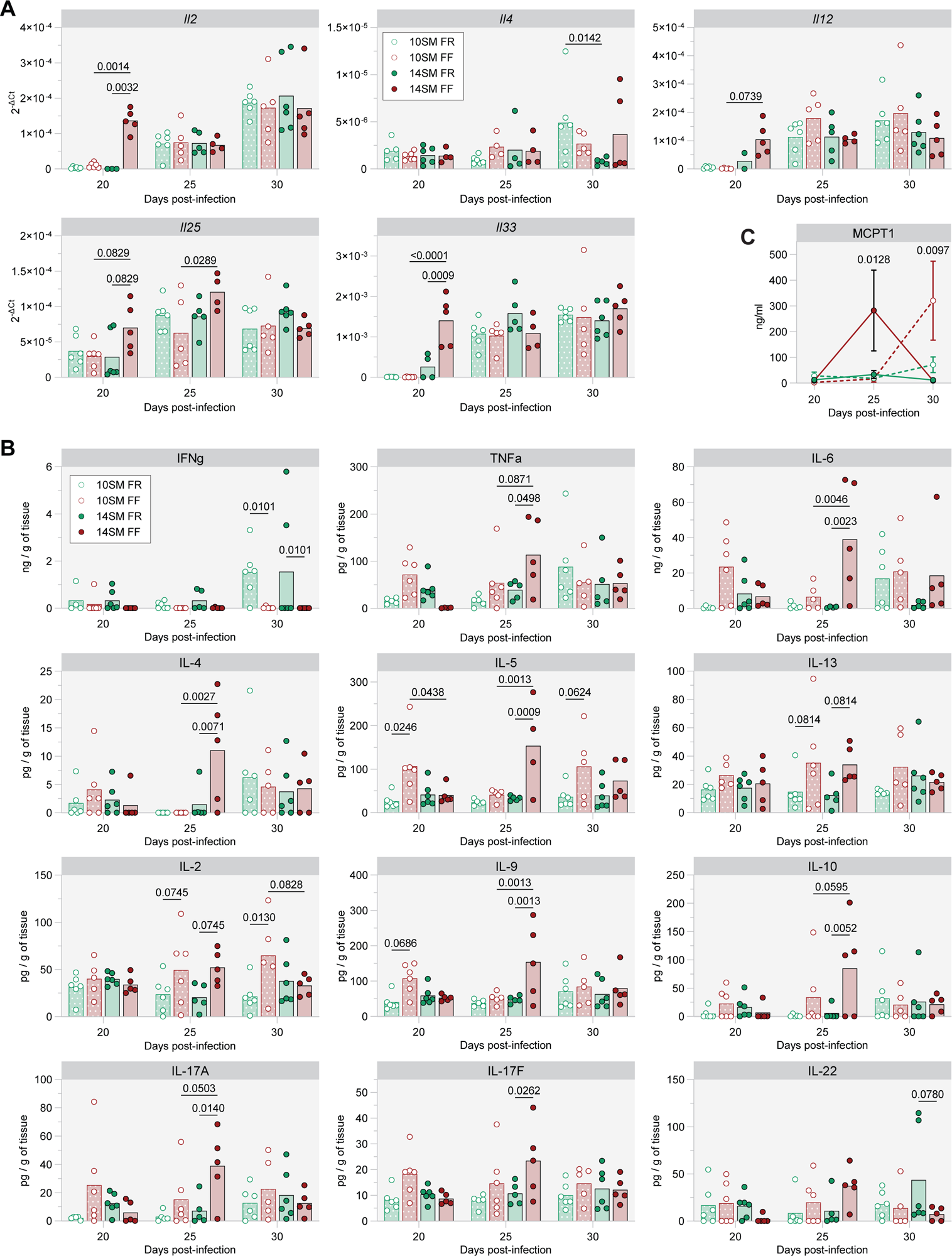

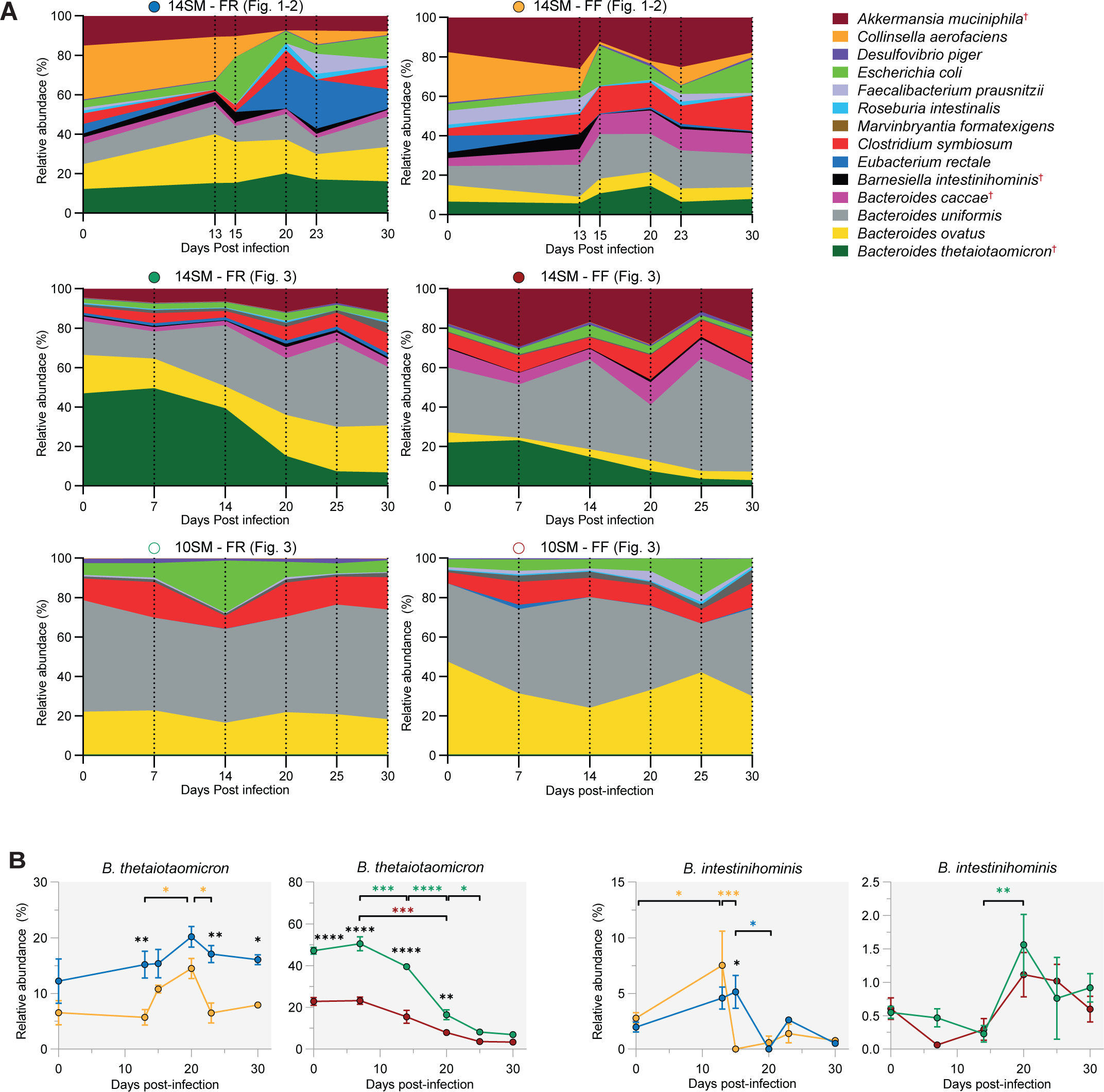

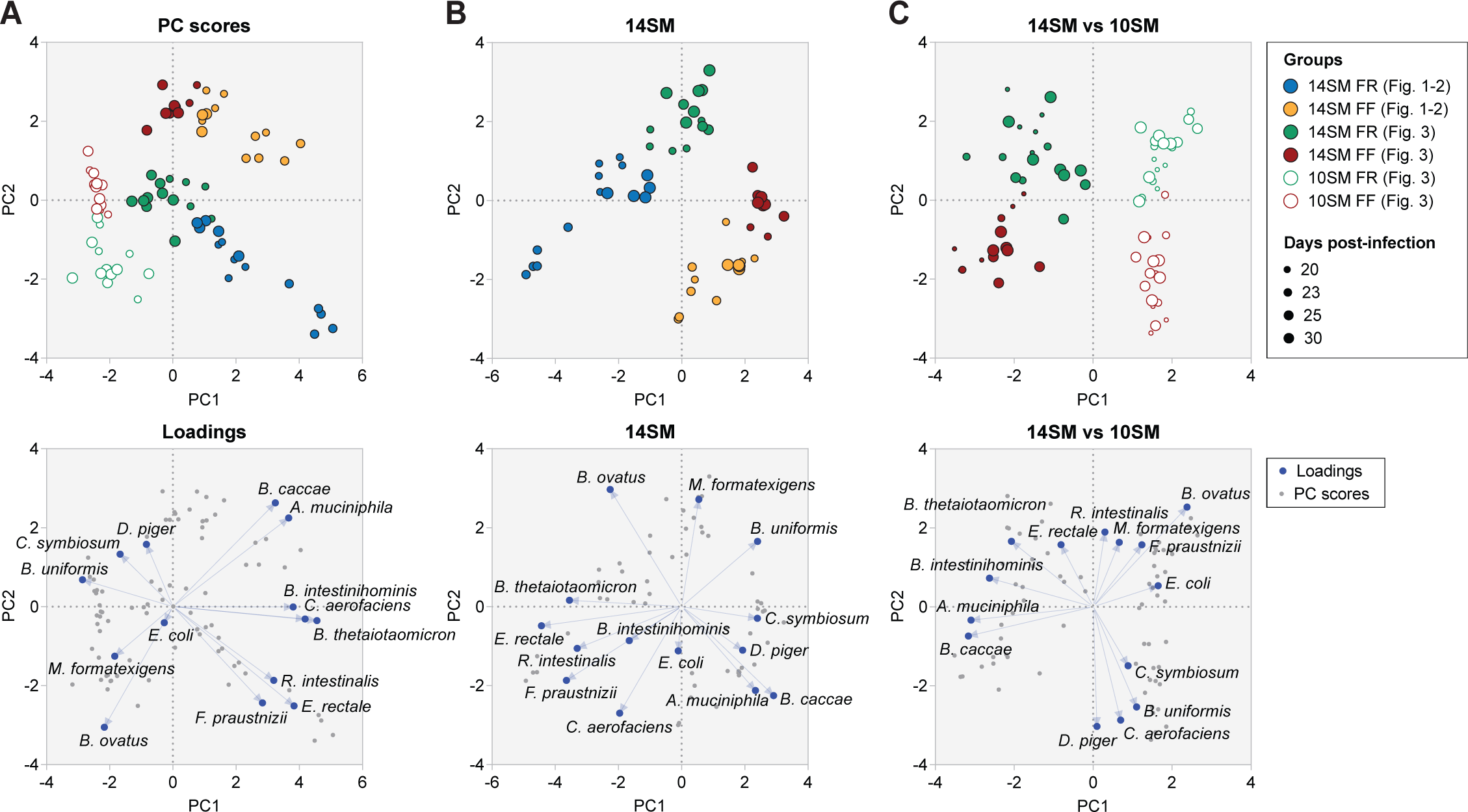

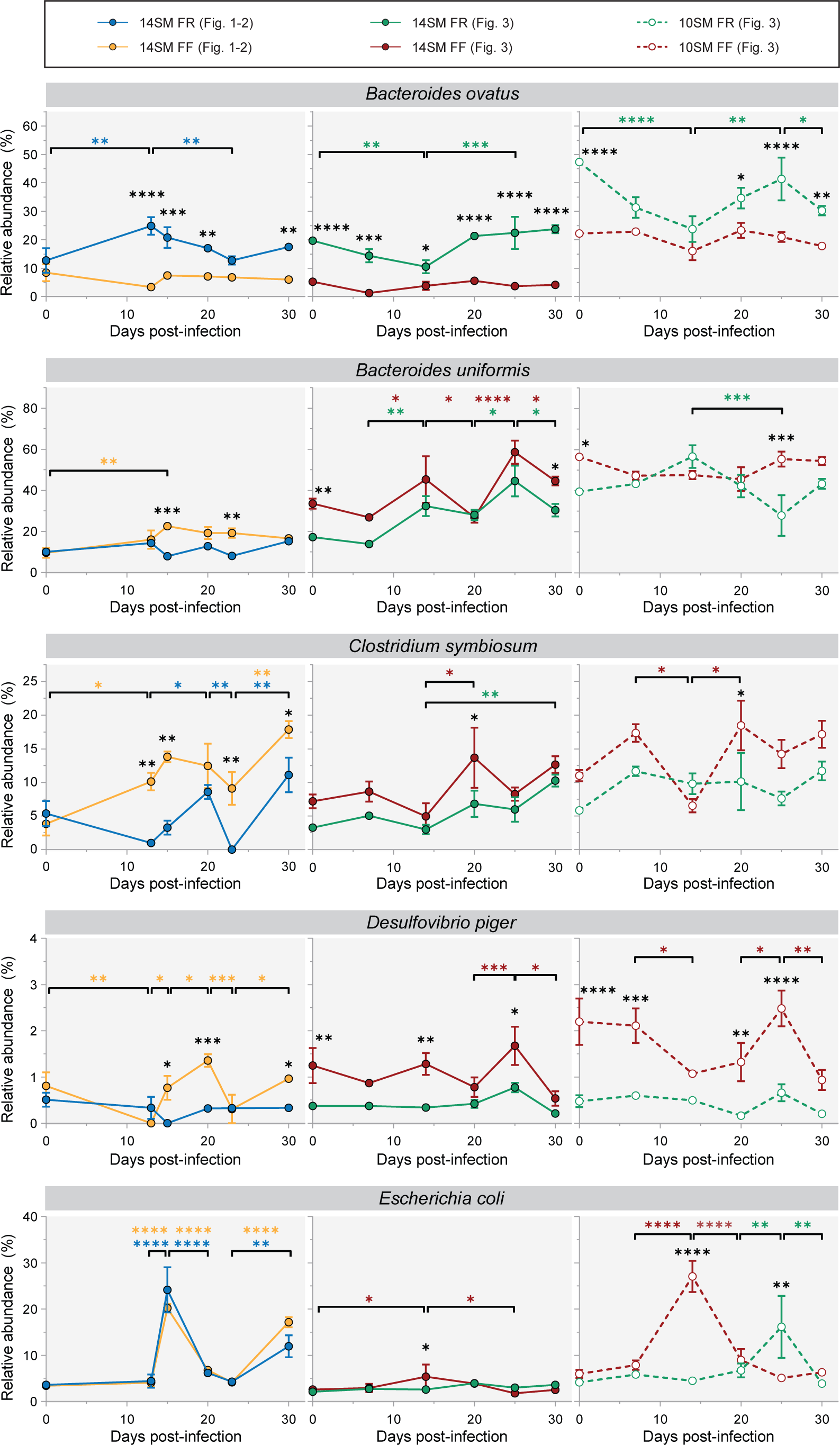

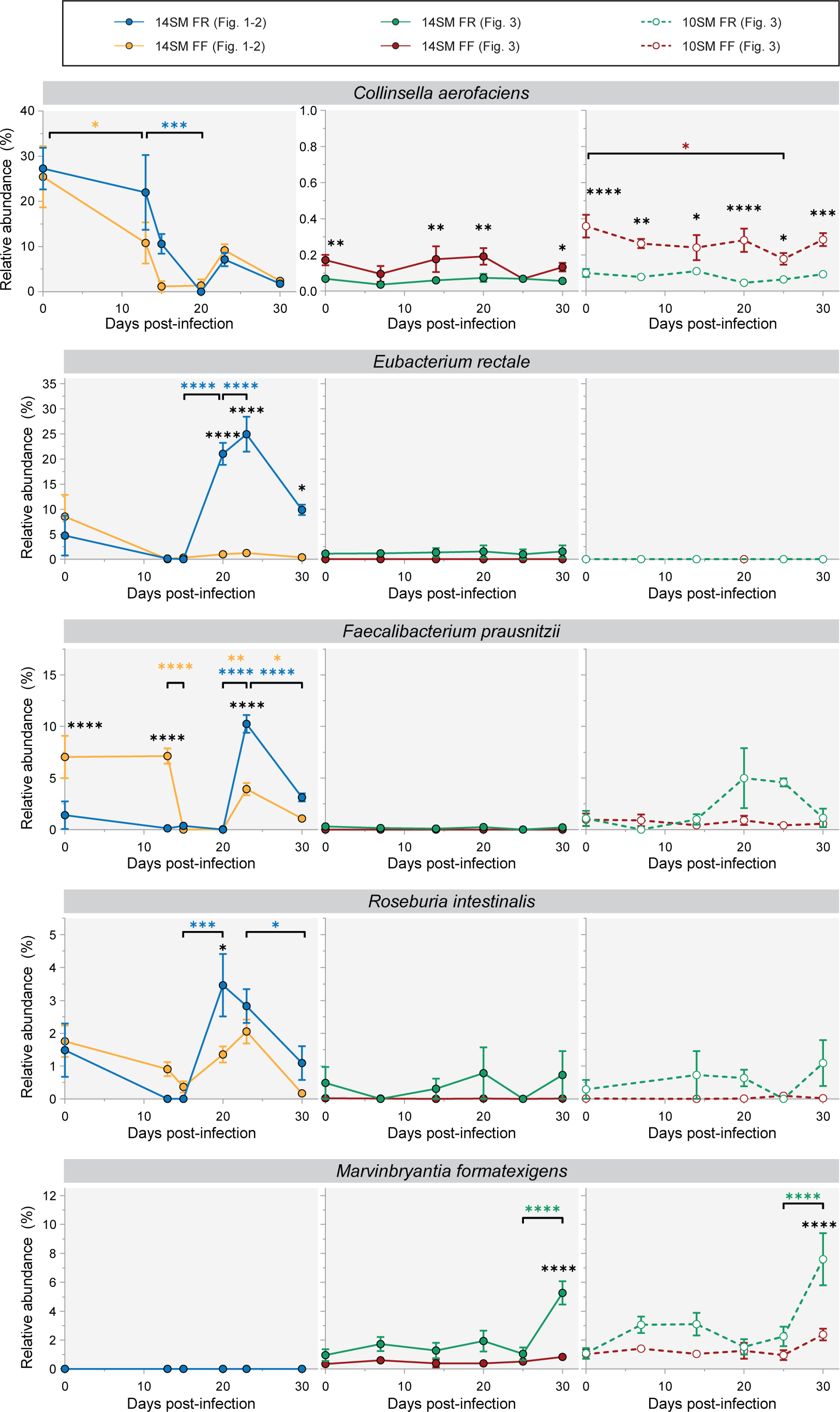

