## Supplementary methods, supplementary results and supplementary figure/table legends for "Mucin-degrading gut bacteria promote anti-parasitic immunity"

### Supplemental methods

##### *Animal diets*

The Fiber-rich diet was a standard autoclaved rodent chow (LabDiet, St. Louis, MO, USA; catalog no. 5013), while the Fiber-free diet was a custom diet manufactured and irradiated by SAFE Diets (Augy, France). The FF diet was manufactured as the previously described TD.140343 diet[1], which is a modified version of the Harlan.TD08810 diet from Envigo (Indianapolis, IN, USA).

##### *14SM and 10SM cultivation and quantification*

All bacterial strains of the 14SM were cultured as previously described[2]. The 10SM consisted of the 14SM without the 4 previously characterized mucin-degrading bacteria, *Akkermansia muciniphila*, *Bacteroides thetaiotaomicron*, *Bacteroides caccae* and *Barnesiella intestinihominis*[1]. The colonization of the 14SM or 10SM was confirmed using strain-specific qPCR primers and relative abundances of the individual bacteria were determined using the same qPCR protocol as described previously[2]. Samples from different dates were run on the same plates in order to avoid plate-specific confounders.

##### *Detection of bacterial glycan-degrading enzyme activities*

The enzymatic activities of α-fucosidase, β-*N*-acetyl-glucosaminidase, and β-glucosidase in fecal samples were determined using *p*-nitrophenyl glycoside-based enzyme assays as described previously[3].

##### *RNA-Seq analysis*

Following adapter removal with Cutadapt[4], reads were mapped and gene counts were generated with STAR 2.7.9a[5]. Transcripts not appearing at least once on average across all samples were removed from subsequent analysis. Samples were normalized using the median of ratios method[6] supported by DESeq2[7] and one outlier was removed based on PC score using the prcomp function of the base R stats 4.0.2 package. Differential expression analysis was performed using DESeq2 1.30.1 with default parameters and $p$-value adjustment using the Benjamini-Hochberg method. Significant genes were identified based on adjusted $p\text{-value} < 0.05$. The expressed log-ratios for significant genes were imported into Ingenuity Pathway Analysis (Qiagen, Hilden, Germany) and analyzed considering all available reference datasets for mice. Pathway analysis was also performed using the enrichGO function of clusterProfiler 3.18.1[8] in R to identify upregulated pathways using Gene Ontology (GO) terms for biological processes. REVIGO[9] was employed to reduce redundancy in GO terms before visualization.

***Serum ELISA for total IgG1 and IgE***

Serum ELISA for total IgG1 and IgE were performed as follows. Plate were coated overnight at room temperature using 0.5 µg/µl rat α mouse IgE purified UNLB (Imtec Diagnostics, 1130-01) or rat α-mouse IgG1 purified UNLB (Imtec Diagnostics, Ardmore, OK, United States; catalog no. 1144-01) capture antibody, respectively, diluted in 0.05 M carbonate/bicarbonate buffer (pH 9.6). Wells were washed three times using a washing buffer consisting of 1% Tween, 154 mM NaCl, 10 mM Trizma Base. 1% w/v of BSA in a TBS buffer (15 mM Trizma-acetate, 136 mM NaCl & 2 mM KCl) was used to block plates by incubating them for 2 hours at room temperature. Afterwards, plates were washed three times using the washing buffer. As controls, Mouse IgG1 Isotype Control UNLB (Imtec Diagnostics, Ardmore, OK, United States; catalog no 0102-01) or Mouse IgE Isotype Control (UNLB), Southern Biotech (Imtec Diagnostics, Ardmore, OK, United States; catalog no 0114-01) was used to generate two-fold serial dilutions. Serum samples were two-fold diluted from 1/40 to 1/1280 using the aforementioned TBS buffer with 0.1% w/v Tween-20 and 1% BSA. Plates were loaded the standards and samples were incubated for 90 minutes at room temperature. Another three wash steps were performed before loading a 1/500 dilution of the 2nd antibody, Goat anti-mouse IgG1-AP, Southern Biotech (Imtec diagnostics, Ardmore, OK, United States; catalog no 1071-04) or Goat anti-mouse IgE-AP, Southern Biotech (Imtec diagnostics, Ardmore, OK, United States; catalog no 1110-04), respectively. The final three wash steps were performed before loading the substrate solution consisting of 0.5 mg/ml phosphate substrate (Sigma-Aldrich, St. Louis, MO, United States; catalog no. S0642-200 TAB) in 1 mM AMP and 0.1 mM $MgCl_2 \cdot 6H_2O$. Plates were incubated for 1 hour at 37°C before they were read at 405 nm using a SpectraMax ABS PLUS spectrophotometer (Molecular Devices, San Jose, CA, United States).

***RNA extraction from mesenteric lymph nodes and colonic tissue***

Mesenteric Lymph nodes (MLN) and colonic tissues were harvested and stored in 1mL of RNAlater™ (Qiagen) in 2mL screw cap tubes and stored at 4°C up to 3 days. After storage, RNAlater was removed and samples were stored at -80°C. For the RNA extraction, a 5mm Autoclaved metal bead (Qiagen) and 1mL of TRIzol™ Reagent (Ambion) was added to the tissue. Homogenization by Bead Beating at full speed was performed followed by a centrifugation at 4°C, 12000 x g for 5min. The clear aqueous phase was transferred in a new safe lock 1.5mL Eppendorf tube and let incubate at RT for 5min before adding 200µl of Pure Chloroform (Fisher BioReagents). Tubes were hand shake for 15sec and let 2-3 min at RT before centrifuged at 12000 x g at 4°C for 15min. Aqueous phase (transparent Upper phase) was transferred in a new safe lock 1.5mL Eppendorf tube and 500µl of Isopropanol was added. Samples were mixed vigoursily for 10sec and let incubate at RT for 10min, followed by a centrifugation at 12000 x g at 4°C for 10min. Supernatant was removed and pellet were resuspended in 1mL of Cold Ethanol 75% followed by a centrifugation at 7500 x g at 4°C for 5min. Supernatant was removed and pellets were dried for 10min at 37°C and resuspended in 50µl of DNase RNase Free water (Thermo Scientific™) and incubated at 56°C for 15min to fully resuspend the pellets. RNA samples were treated with DNase1 (Invitrogen), according to manufactor instructions and Purified using RNeasy Mini (Qiagen) according to manufactor Clean Up protocol. Purified RNA samples were eluted in 30µl of DNase RNase free Water twice (by transferring the eluated product back in the column) before been quantify with Nanodrop.

***RNA extraction from Colon Content Samples***

2 Pellets of colon Content were stored in 500µl of RNAProtect™ (Qiagen) in 2mL screw cap tubes and stored at 4°C for up to 3 days. After storage, samples were centrifuged at 10000 x g for 10min and RNAProtect™ was removed and pellets were stored at -80°C. For the RNA extraction, 250µl of ceramic beads (Macherey Nagel) were added to the pellets with 500µl of Buffer A (200mM NaCl, 200mM Tris, 20mM EDTA pH8.0), 210µl of SDS 20% (Carl Roth) and 500µl of Phenol: Chloroform (125:24:1) pH4.3 (Fisher BioReagents). Samples were homogenized by Bead beating for 5min and centrifuged at 18000 x g for 3min at 4°C. Aqueous phase was recovered and 500µl of Phenol: Chloroform (125:24:1) pH4.3 were added and mixed by gentle inversion. Samples were centrifuge at 4°C at 18000 x g for 3min and aqueous phase was mixed with 1/10 volume of 3M Sodium acetate pH5.5 and 1 volume of Cold 100% Ethanol, followed by an incubation at -20°C for 20min. Samples were centrifuged at 4°C for 20min at 18000 x g and supernatant was discarded to resuspend pellet in 500µl of Cold Ethanol 70%. Centrifugation at 4°C for 5min at 18000 x g and washing with Ethanol 70% was repeated. Pellet was air dried and resupended in 50µl of DNase RNase Free water. RNA samples were treated with DNase1 (Invitrogen), according to manufactor instructions and Purified using RNeasy Mini (Qiagen) according to manufactor Clean Up protocol. Purified RNA samples were eluted in 30µl of DNase RNase free Water twice (by transferring the eluated product back in the column) before been quantify with Nanodrop.

***RT-qPCR***

Targeted analysis of cytokine and transcription factor expression was determined by RT-qPCR of RNA extracted from mesenteric lymph nodes (MLN) and colonic tissue (colon). The cDNA library was prepared by combining 500 µM dNTP Set (100 mM) Solution (Invitrogen, Waltham, USA), 2.5 µM Random Primers (Invitrogen), 100 ng/µl of RNA sample and 1µl of ddH2O per sample, followed by heating to 65°C for 5 min and incubation on ice for at least 1 min. Next, 1× SSIV Buffer, 100 mM DTT, 40 U RNaseOUT (Invitrogen, Waltham, USA), Recombinant Ribonuclease Inhibitor (Invitrogen, Waltham, USA), and 200 U Superscript IV Reverse Transcriptase (Invitrogen, Waltham, USA) was added to each sample and incubated at 23°C for 10 min. Samples were incubated at 50°C for 10 min, then heat-inactivated at 80°C for 10 min.

For qPCR, a master mix consisting of 1× Buffer, $MgCl_2$ (concentration in mix differs depending on the primers used, see **Table S4**), 400uM dNTP, 1× SYBR Green I Nucleic Acid Gel Stain, 10,000× concentrate in DMSO (Invitrogen, Waltham, USA), and 0.5 U Platinum *Taq* DNA Polymerase (Life Technologies, Carlsbad, USA), was added to 1 µl each cDNA sample along with forward and reverse primers (the concentrations of specific primers are listed in **Table S4)**. The qPCR cycle for primers targeting Transcription factors and cytokines consisted of pre-denaturation at 94°C for 5 min, followed by 40 cycles of 20 sec denaturation at 94°C, 50 sec annealing at 60°C, and 45 sec extension at 72°C. Samples were held at 72°C for 5 min post-extension and then a melting curve was generated by heating from 65°C to 95°C with 0.3°C interval increases over 15 sec. For primers targeting *C1galt1* and *Galnt3* a different qPCR program was used: pre-denaturation at 95°C for 2 min, followed by 45 cycles of 15 sec denaturation at 95°C, 15 sec annealing at 60°C, and 45 sec extension at 72°C.

##### *Goblet cell counting*

Goblet cell counting was performed on a small piece (~0.5 cm) of methacarn-fixed and Alcian blue-stained cecal tissue. To preserve the mucus within the intestinal samples, a modified direct pretreatment in 99% ethanol and toluene was performed, omitting the usual previous ascending ethanol steps before embedding the samples in paraffin. All samples were cut using a microtome at 3 µm thickness before performing automated HE (Tissue-Tek Prisma Plus and Film, Sakura, Alphen aan den Rijn, Netherlands) and Alcian blue (Artisan Link Pro Special Staining System, Dako, Glostrup, Denmark) staining according to the manufacturers' instructions. The numbers of goblet cells per crypt were counted and averaged for each individual mouse before the mean per experimental group was determined. The researcher, S.T., who counted the goblet cells, was blinded for both the individual time points and dietary groups.

#### Supplementary results

##### ***Infection with a high egg number leads to chronic infection in 14SM-colonized mice***

Infection of *T. muris* with a high egg number (~450) in conventional or specific-pathogen-free (SPF) mice, results in an acute infection, and only lower number of eggs (<40) result in a chronic infection[10]. In our synthetic human gut microbiota under standard dietary conditions (that is fed the FR diet), infection with a high number of eggs caused a low level infection resulting in a chronic infection. The reasons for this could be rooted in the fact that our study uses a synthetic human gut microbiota, as opposed to the complete, native murine microbiota that may be more suitable for egg hatching of the naturally occurring murine whipworm or improving the ability of the larvae to establish in the gut. Considering that this a new model, it is particularly an important point because previous studies have shown that the gut bacteria are essential for effective egg hatching, as the eggs do not hatch in the absence of gut bacteria *in vitro* or in germ-free mice[11,12].

##### **T. muris** ***infection impairs 14SM equilibrium***

The 14SM microbiota composition was different between the first experiment (**Fig. 1, 2**) and the second experiment (**Fig. 3**) at the time of infection, with more *Collinsella aerofaciens* and *Feacalibacterium prausnitzii* in the first experiment and more *Bacteroides thetaiotaomicron* in the second experiment (**Fig. S4A**). These 2 experiments were performed independently with mice of different ages (22 weeks in **Figs. 1, 2** versus 8–12 weeks in **Fig. 3**). However, the experiment performed for Figure 3 was also performed in 3 independent batches of colonization, but shows consistency between batches. Thus, it is more likely that the difference of microbiota composition between the 2 experiments is linked to the age of the mice used in each experiment. Although the composition of the microbiota evolved during the infection, Principal Component Analysis (PCA) revealed that distinct compositions were maintained over time between experiments and between experimental groups within each experiments (**Fig. S5A**). Among 14SM-colonized groups, FF-fed groups were characterized by a higher proportion of 2 mucin-degrading bacteria *A. muciniphila* and *B. caccae*, in addition to *D. piger* and *C. symbiosum*, while the FR-fed groups were characterized by a higher proportion of *B. ovatus* (**Fig. S5B**). As expected, the main factors discriminating between 10SM and 14SM microbiota were the 4 mucin-degrading bacteria *A. muciniphila*, *B. caccae, Bacteroides thetaiotaomicron* and *B. intestinihominis* (**Fig. S5C**).

#### Supplementary figure legends

**Figure S1. Dietary fibers promote Th1 responses during *T. muris* infection.**

(A) Weekly assessed weights of *T. muris* infected mice. Error bars represent SEM (n = 4–23); ****$p<0.0001$ between FR and FF group as determined by a 2-way ANOVA and adjusted with the FDR method of Benjamini and Hochberg.

(B, C) Analysis of data generated by RNA-seq of RNA extracted from colonic tissue at 30 dpi (n = 4). (B) Graphical summary generated by IPA showing the major upregulated genes and pathways as well as their interactions. (C) Relative gene expression (FR/FF) of the 5 most significant activated pathways in the colonic tissue under FR conditions. Pathways are based on GO terms for biological processes and REVIGO was used to reduce redundant terms.

(D) Relative expression of indicated transcripts in mesenteric lymph nodes and colonic tissue (colon) over time after T. muris infection. Error bars represent SEM (n = 3–5); *p* values indicate comparisons between FR and FF groups as determined by a 2-way ANOVA and adjusted with the FDR method of Benjamini and Hochberg.

(E) Total IgG1 and total IgE concentrations in mouse serum (n = 3–5).

**Figure S2. Dietary fibers and mucin-degrading bacteria regulate the immune cell profile during worm infection.**

(A) Maturation stage of larvae counted in panel A. Each dot represent the average estimation of all larvae per mouse (n = 5–6). Multiple Mann-Whitney comparisons and *p* values adjusted with the FDR method of Benjamini and Hochberg.

(B) Fecal Lipocalin-2 (LCN-2) levels assessed by ELISA over the course of *T. muris* infection. Error bars represent SEM (n = 5–6).

(C) Proportion of indicated immune cell populations in the colonic lamina propria (n = 4–6). Two-way ANOVA and *p* values adjusted with the FDR method of Benjamini and Hochberg.

**Figure S3. Dietary fibers and mucin-degrading bacteria regulate the cytokine response during worm infection.**

(A) Relative expression of indicated transcripts in colonic tissue (n = 2–6).

(B) Secreted cytokines in colon tissue (n = 4–6).

(C) Serum mast cell protease 1 (MCPT1) concentrations. Error bars represent SEM (n = 5–6). Two-way ANOVA and *p* values adjusted with the FDR method of Benjamini and Hochberg.

**Figure S4. Evolution of synthetic microbiota compositions during *T. muris* infection.**

(A) Microbial composition of the gut microbiota at indicated time points after *T. muris* infection (denoted by dotted lines). Relative bacterial abundances were determined by qPCR of DNA extracted from fecal pellets using primers specific to each bacterial strain. Red crosses denote known mucin-degrading bacteria.

(B) Relative abundances of *B. thetaiotaomicron* and *B. intestinihominis*. Error bars represent SEM (n = 2–6); two-way ANOVA and multiple comparisons with the FDR method of Benjamini and Hochberg. Significance labels indicate comparisons between FR versus FF groups (black), or between indicated timepoints among FR-fed mice of Fig. 1–2 (blue) FF-fed mice of Fig. 1–2 (yellow), FR-fed mice of Fig. 3 (green) and FF-fed mice of Fig. 3 (red).

**Figure S5. Principal Component Analysis (PCA) of microbiota composition**

PCA of microbiota composition among all groups (A, n = 81), 14SM-colonized groups (B, n = 52) and Fig. 3-related groups (C, n = 71). The upper line shows the PC scores of microbiotas and the bottom line shows the weight/loading of each strain in the PCA.

**Figure S6. Relative abundances of *B. ovatus*, *B. uniformis*, *C symbiosum*, *D. piger* and *E. coli* during *T. muris* infection.**

Error bars represent SEM (n = 2–6); two-way ANOVA and multiple comparisons with the FDR method of Benjamini and Hochberg. Significance labels indicate comparisons between FR versus FF groups (black), or between indicated timepoints among FR-fed mice of Fig. 1–2 (blue) FF-fed mice of Fig. 1–2 (yellow), FR-fed mice of Fig. 3 (green) and FF-fed mice of Fig. 3 (red).

**Figure S7. Relative abundances of *C. aerofaciens*, *E. rectale*, *F. prausnitzii*, *R. intestinalis* and *M. formatexigens* during *T. muris* infection.**

Error bars represent SEM (n = 2–6); two-way ANOVA and multiple comparisons with the FDR method of Benjamini and Hochberg. Significance labels indicate comparisons between FR versus FF groups (black), or between indicated timepoints among FR-fed mice of Fig. 1–2 (blue) FF-fed mice of Fig. 1–2 (yellow), FR-fed mice of Fig. 3 (green) and FF-fed mice of Fig. 3 (red).

#### Supplementary table legends

**Table S1.** Relative gene expression (FR/FF) based differential expression analysis using DESeq2.

**Table S2.** Significantly upregulated pathways in FR diet based on gene oncology (GO) terms for biological processes.

**Table S3.** Significantly upregulated pathways in FF diet based on gene oncology (GO) terms for biological processes.

**Table S4.** Primers used for RT-qPCR.

**Table S5.** Correlations between the fucosidase activity and the relative abundance of 14SM members.

**Table S6.** Correlations between the N-acetyl-glucosaminidase activity and the relative abundance of 14SM members.

**Table S7.** Correlations between the Glucosidase activity and the relative abundance of 14SM members.
